# The ammonia oxidizing bacterium *Nitrosomonas eutropha* D23 blocks T helper 2 cell polarization via IL-10 – mediated interference with dendritic cell activation

**DOI:** 10.1101/2020.06.30.175414

**Authors:** Damien Maura, Nazik Elmekki, Alex Goddard

## Abstract

The prevalence of atopic diseases has been steadily increasing since the mid 20^th^ century, a rise that has been linked to modern hygienic lifestyles that limit exposure to microbes and immune system maturation. Overactive type 2 CD4+ helper T (Th2) cells are known to be closely associated with atopy and represent a key target for treatment. In this study, we report that the ammonia oxidizing bacteria (AOB) *Nitrosomonas eutropha* D23, an environmental microbe that is not associated with human pathology, effectively suppresses the polarization of Th2 cells and production of Th2-associated cytokines (IL-5, IL-13, & IL-4) by human peripheral blood mononuclear cells (PBMC). We show that AOB inhibit Th2 cell polarization not through Th1-mediated suppression, but rather through an IL-10-mediated mechanism that interferes with the activation of dendritic cells, as evidenced by a reduction in Major Histocompatibility Complex Class II (MHC II) and CD86 expression following AOB treatment. This is the first report of immunomodulatory properties of AOB and supports the development of AOB as a potential therapeutic for atopic diseases.

**GRAPHICAL ABSTRACT:** 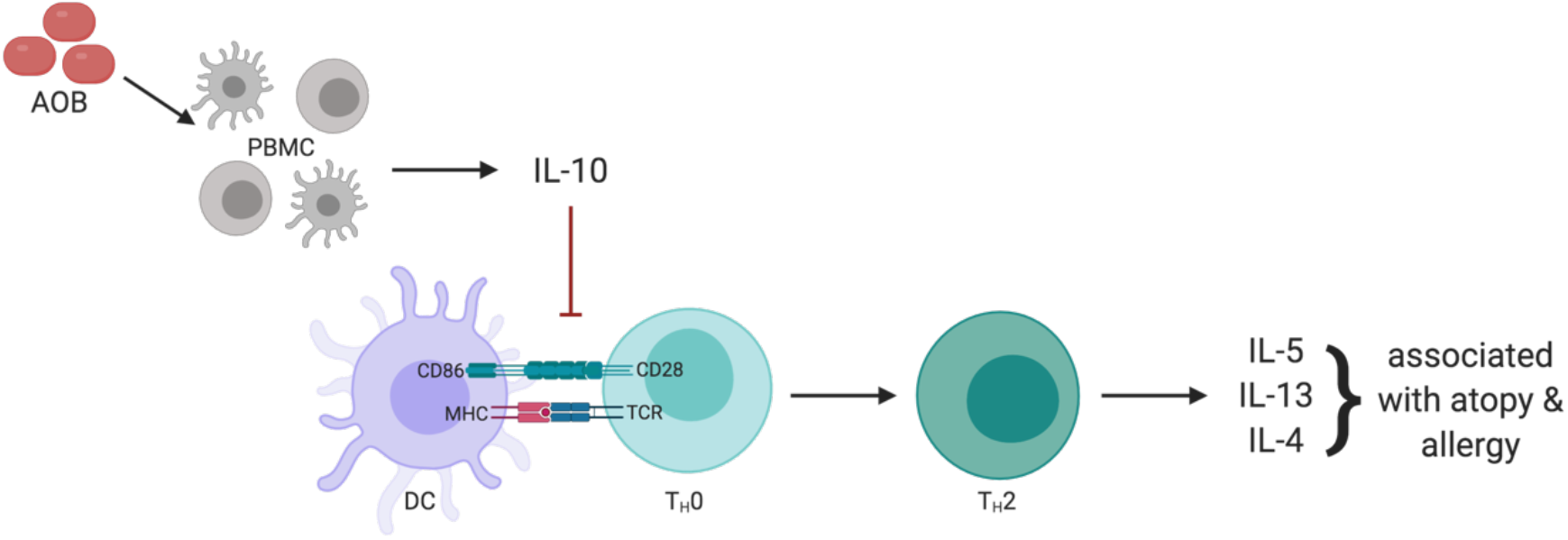

*Created with BioRender.com*

## INTRODUCTION

Since the mid 20^th^ century, the prevalence of atopic diseases has been steadily increasing with more than 200 million people worldwide suffering from diseases such as asthma, allergic rhinitis, or food allergies ^1^. The rapid increase in atopy prevalence cannot be explained by changes in population genomics. Rather, the association with westernized societies suggests that environmental factors such as diet, antibiotic exposure, and other modern hygienic lifestyles play a crucial role in the etiology of these conditions ^2–5^.

Notably, reduced exposure to non-commensal microbes, particularly early in life, as well as a loss of biodiversity within the human microbiome has been associated with atopy ^5–8^. Exposure to microbes is essential for immune system development and training. Microbes can reinforce barrier immunity and condition innate cells to promote effector B and T cell responses against pathogens, and more specifically balance the relative numbers and specificity of different CD4+ T helper cell populations (Th1, Th2 and Th17) and regulatory T cells (Tregs) ^9–11^. When underexposed to foreign antigens, the immune system is not trained to react appropriately to non-noxious antigenic stimuli, which leads to disproportionate responses to self-tissues (auto-immune diseases) or external triggers that are otherwise innocuous stimuli (atopic diseases) ^12,13^.

Atopic diseases are marked by an exaggerated, systemic type 2 inflammatory reaction to otherwise innocuous stimuli that induce high levels of immunoglobulin E (IgE) production by B cells, eosinophilia, and various associated inflammatory immune responses ^14^. Therapeutic approaches that interfere with Th2 cells and their cytokine mediators such as IL-5, IL-13, & IL-4 have shown promise, highlighting the potential of targeting this pathway ^14^.

Given the role of microbes in immune system maturation, several studies have focused on identifying bacteria that have the ability to modulate the Th2 pathway and improve allergy symptoms in preclinical models ^6^. Probiotic or commensal bacteria such as *Lactobacillus*, *Weissela cibaria* WIKIM28 or *Bacteroides fragilis* were found to interfere with the Th2 pathway by inducing Th1 cells or Tregs ^15–18^. Exposure to agrarian conditions is also suggested to be protective, leading investigators to search for microbes in cowsheds and other farming environments. Cowshed bacteria, such as *Acinetobacter lwoffii* F78, *Lactococcus lactis* G121, and *Staphylococcus sciuri* W620, were shown to shift T cell polarization towards a Th1 response via dendritic cell conditioning ^6,19–21^. These cowshed bacteria highlight the potential to identify novel immuno-modulatory bacteria from farm environments.

Farm soils frequently contain high concentrations of ammonium due to animal waste and fertilizers, and are consequently niches containing ammonia oxidizing bacteria (AOB) ^22,23^. These soil chemolithoautotrophic Gram negative bacteria play a critical role in the global nitrogen cycle ^22^. They can extract energy by oxidizing ammonia, a process generating the immunomodulatory molecule nitric oxide as a byproduct ^24,25^. Interestingly, AOB were detected in human microbiomes and have been hypothesized to be depleted in recent decades due to modern hygienic lifestyles ^26–28^. Therefore, we postulated that AOB could play a role in the human microbiome’s ability to regulate the immune response.

In this study, we present the first report that the AOB *Nitrosomonas eutropha* D23 inhibit Th2 immune polarization in human primary immune cells. Unlike many other Th2 suppressing bacteria, AOB do not to require the classical Th1 pathway to achieve this Th2 suppression, but rather dampen dendritic cells’ ability to activate Th2 cells in a process that involves IL-10.

## RESULTS

### AOB inhibit Th2 immune polarization in human PBMC

To test the effect of AOB on immune responses, we developed a co-culture model with human peripheral blood mononuclear cells (PBMC) derived from healthy subjects. While AOB would not directly encounter blood and its constituents frequently, PBMC provide a well-studied model to test host-microbe interactions, and allow for investigations that are inclusive of many immune cell-types and potential signaling pathways.

First, we validated this host-microbe model by investigating AOB toxicity to PBMC and *vice versa*. AOB were added to PBMC at a ratio of four bacteria per PBMC for 72 hours. AOB did not significantly affect PBMC cell viability as measured by the metabolic activity dye WST-1 or LIVE/DEAD staining (Figure S1a-b, p>0.05). Furthermore, PBMC did not adversely affect AOB viability, as nitrite production, a measure of AOB activity, remained unchanged in presence or absence of PBMC (Figure S1c p>0.05). Moreover, the ability of AOB, but not PBMC, to produce nitrite in RPMI media suggests the bacterial cells are active, metabolizing ammonia present in the media that is potentially produced by PBMC metabolism (Figure S1c). Thus, AOB and PBMC can effectively be co-cultured and their respective activities assessed.

We then tested if AOB modulate Th2 immune response pathways associated with atopic diseases ^29^. We used PBMC from a healthy, non-atopic human donor, and treated them with exogenous factors to induce a Th2-polarized phenotype. First, we stimulated PBMC for 3 days using a proprietary Th2 differentiation kit (R&D Systems) and measured cytokine production in the culture supernatant by ELISA. Th2 stimulation led to the production of 826 ± 253 pg/mL IL-5 and 495 ± 177 pg/mL IL-13, two key markers of Th2 polarization. In contrast, unstimulated cells did not produce any detectable levels of these cytokines (Figure 1a-b).

**Figure 1:**
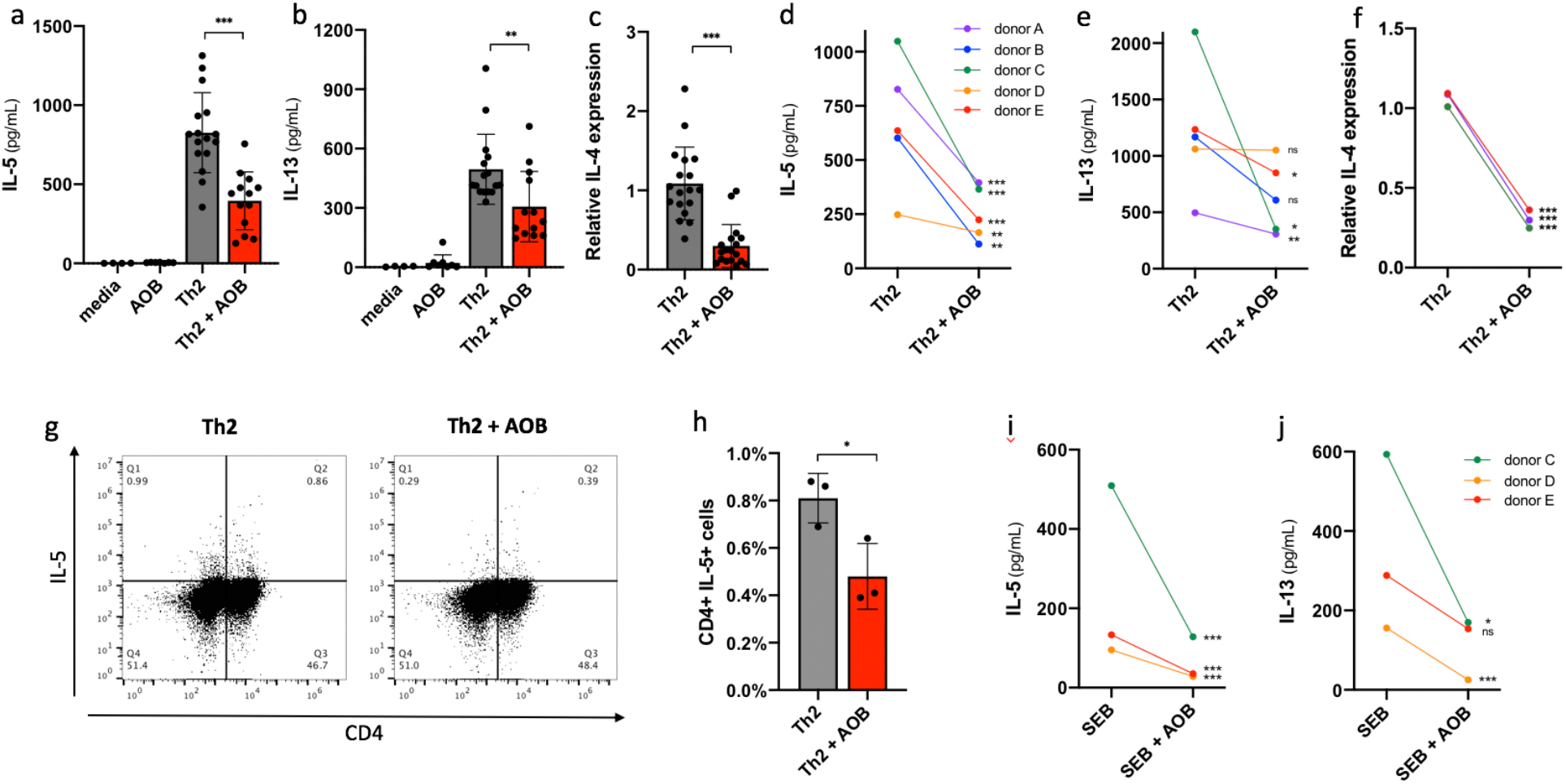
AOB inhibit Th2 immune polarization in human PBMC. (a-f) IL-5 and IL-13 supernatant levels are reduced with AOB pretreatment of PBMC prior to Th2 differentiation cocktail stimulation; measured by ELISA of supernatants 72h post-stimulation. IL-4 gene expression is also reduced with AOB pretreatment; measured by qPCR 72h post-stimulation. AOB pre-treatment prior to Th2 differentiation cocktail stimulation of PBMCs from donor A reduced the levels of IL-5 (a) & IL-13 (b) as well as IL-4 (c) gene expression (n**≥**12, one-way ANOVA with multiple comparisons or unpaired t-test). (d-f) IL-5 (d), IL-13 (e), and IL-4 (f) levels/expression are reduced in AOB-pretreated PBMC from 3-5 different donors (n**≥**6 replicates per donor, shown on the figure is unpaired t-test p value for individual donors, unpaired t-test p value for aggregated donors: p<0.0001 for IL-5, p<0.01 for IL-13, p<0.001 for IL-4). (g, h) The number of CD4+ IL-5+ cells is reduced with AOB pretreatment of PBMC following Th2 differentiation cocktail stimulation; measured by flow cytometry with anti-CD4 PerCP cy5.5 and anti-IL-5 PE antibodies applied 72 hours post-stimulation. (g) Representative dot plot from donor C cells showing the reduction in CD4+ and IL-5+ cells. (h) The percentage of Th2 cells as CD4+ IL-5+ are reduced with AOB pretreatment (n=3, donor C, unpaired t-test). IL-5 (i) and IL-13 (j) supernatant levels are reduced in the presence of AOB 7 days post-stimulation with SEB (1 μg/mL), measured by ELISA (n=3 per donor, 3 donors, unpaired t-test, p<0.0001 for IL-5 and IL-13 for aggregated donor data sets).

We next tested if AOB modulate the Th2 polarization phenotype. PBMC from the same healthy donor were incubated with AOB one hour prior to Th2 stimulation, and cytokines were measured by ELISA in culture supernatants 72 hours after Th2 stimulation. The addition of AOB significantly reduced IL-5 and IL-13 production as compared to the samples pre-treated with vehicle (Figures 1a-b, p<0.01). Similarly, AOB treatment reduced the gene expression of IL-4, another key Th2 marker (Figure 2c, p<0.001); see Methods for comment on IL-4 assessments. As PBMC from different donors can model diversity in the population, we tested the effect of AOB on Th2 cytokine production from PBMC derived from four other donors (Figures 1d-e). AOB pretreatment significantly reduced IL-5 production in 5 of 5 donors tested (Figure 1d, p<0.01) and IL-13 production in 3 of 5 donors (Figure 1e, p<0.05). AOB also reduced IL-4 expression in 3 of 3 donors tested (Figure 1f, p<0.001). The IL-5 and IL-13 inhibition was also observed 120 hours following stimulation, suggesting temporal stability of the inhibition (Figure S2 a-b, p<0.05). As we saw robust modulation of IL-5 across donors, we used IL-5 as a main readout for the rest of the study.

**Figure 2:**
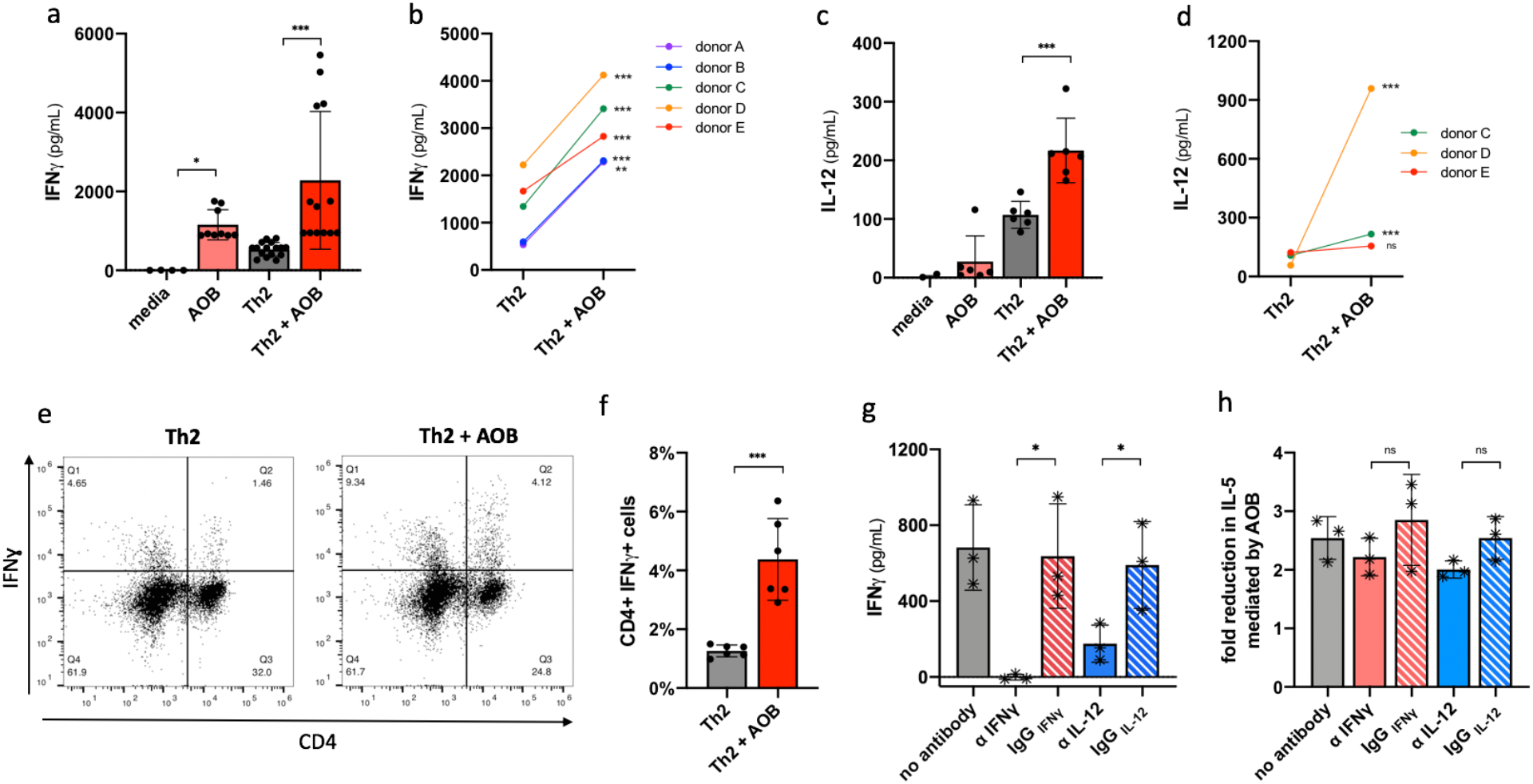
AOB induce Th1 polarization but Th1 are not necessary for Th2 inhibition. (a-d) IFNγ (a, b) and IL-12p70 (c, d) supernatant levels are increased with AOB pretreatment of PBMC prior to Th2 kit stimulation; measured by ELISA. IFNγ (a) and IL-12p70 (c) levels are increased in PBMC from donor A (n=9, one-way ANOVA with multiple comparisons). IFNγ (b) is induced in AOB pre-treated PBMC from 5 donors and IL-12p70 (d) is induced in PBMC from 2 of 3 donors tested (n**≥**6 per donor, unpaired t-test, p<0.0001 for IFNγ and p<0.0015 for IL-12 aggregated donor datasets). (e, f) The number of CD4+ IFNγ+ cells is increased with AOB pretreatment of PBMC prior to Th2 differentiation cocktail stimulation as measured by staining with anti-CD4 PerCP cy5.5 and anti-IFNγ FITC antibodies 72 hours post-stimulation. (e) Representative dot plot from donor C PBMC showing an increase in the percentage of CD4+ and IFNγ+ cells. (f) Percentage of Th1 cells as CD4+ IFNγ+ are increased with AOB pretreatment (n=6, unpaired t-test). (g, h) Addition of IFNγ or IL-12 neutralizing antibodies to PBMC prior to AOB pretreatment does not prevent AOB inhibition of Th2. IFNγ (g) production is reduced in presence of 10μg/mL IFNγ or IL-12 neutralizing antibodies in PBMC from 3 donors (n=3 per donor, one point per donor, one-way ANOVA with multiple comparisons). AOB-mediated reduction in IL-5 production (h) is unaffected in presence of 10μg/mL IFNγ or IL-12 neutralizing antibodies in PBMC from 3 donors (n=3 per donor, one point per donor, one-way ANOVA with multiple comparisons).

IL-5 can be generated from multiple sources, including CD4+ T cells and eosinophils. We thus explored whether AOB were specifically inhibiting IL-5 production in CD4+ T cells that would induce an atopic response. We immunostained Th2 stimulated PBMC with fluorescently labeled antibodies against CD4 and IL-5, and then quantified the proportion of double-positive cells with and without AOB pretreatment using flow cytometry. AOB pretreatment significantly reduced the percentage of CD4+ IL-5+ cells following Th2 stimulation (Figures 1g-h, p<0.05), suggesting AOB can prevent the production of Th2 associated cytokines from CD4+ T cells.

Finally, we sought to confirm the ability of AOB to mitigate Th2 polarization using a different Th2 stimulation methodology. We used Staphylococcal enterotoxin B (SEB), a potent inducer of Th2 response, and a major contributor to the pathology of atopic dermatitis ^30^. We confirmed that SEB was able to induce IL-5 and IL-13 production in 3 of 3 donors (Figure 1i-j). Furthermore, AOB pretreatment of PBMC one hour prior to SEB stimulation also significantly reduced IL-5 and IL-13 production in multiple donors (Figures 2i-j, p<0.05). Overall, these data, using multiple stimulation paradigms and multiple donors, demonstrate that AOB can suppress the induction of the human Th2 response.

### AOB – mediated Th2 inhibition is largely independent of Th1 signaling

Having established that AOB can suppress the induction of Th2 signaling, we sought to understand what upstream mechanisms could contribute to this suppression. Th1 cells can block Th2 polarization via the effector molecule IFNγ, a mechanism used by *Lactobacillus* to interfere with Th2 signaling ^17^. Therefore, we first tested if AOB induce the production of IFNγ. AOB pretreatment significantly induced IFNγ production, both in unstimulated and in Th2 stimulated PBMC (by Th2 differentiation kit) (Figure 2a, p<0.001). This observation was replicated in PBMC from all 5 donors tested (Figure 2b, p<0.01).

Th1 polarization involves the dendritic cell – mediated release of IL-12, which in turn leads to IFNγ release from CD4+ T cells ^31^. Therefore, we tested if AOB pretreatment specifically induced a Th1 phenotype by measuring AOB impact on i) the DC-derived Th1-inducing cytokine IL-12 and ii) IFNγ production by CD4+ T cells specifically. As shown in Figure 2c, AOB pretreatment increased the levels of IL-12 in culture supernatants following a Th2 stimulus (p<0.001), an observation validated in 2 of 3 donors (Figure 2d). To determine if IFNγ was being produced by CD4+ T cells specifically, PBMC were immunostained with fluorescently labeled antibodies against CD4 and IFNγ and analyzed using flow cytometry. We observed a 3.5-fold increase in the proportion of CD4+ IFNγ+ cells after AOB treatment (Figures 2e-f, p<0.001). Altogether, these data strongly suggest that AOB induce a Th1 response.

Next, we investigated whether Th1 signaling was necessary for the AOB-mediated suppression of Th2 responses. To remove the influence of Th1 signaling, we applied either IFNγ or IL-12 neutralizing antibodies to the PBMC prior to AOB treatment and Th2 stimulation. Both IFNγ or IL-12 neutralizing antibodies drastically reduced the levels of IFNγ in the supernatant as compared to isotype controls, thus validating that Th1 pathway output and development are suppressed, respectively (Figure 2g, p<0.05). However, neither neutralizing antibody treatment significantly influenced the AOB-mediated suppression of IL-5 (Figure 2h, p>0.05). In sum, these data demonstrate that while Th1 responses are induced by AOB treatment, the resulting Th1 responses and IFNγ production are not major contributors on AOB-mediated inhibition of Th2 polarization.

### Th2 inhibition by AOB involves IL-10 – mediated suppression of dendritic cell activity

The ability of microbes to modulate the interaction between dendritic cells & T cells has been well established ^32–34^. Cowshed bacteria can interfere with the ability of dendritic cells to promote Th2 polarization by blocking the expression of the Notch ligand Jagged-1 ^35^. However, AOB did not reduce the expression of Jagged-1 (*data not shown*), suggesting a different mechanism is involved. Dendritic cells activate CD4+ T cells through antigen presentation via MHC II and co-stimulation factors such as CD86/CD80 ^36,37^. Therefore, we tested if AOB modulate dendritic cell activation by downregulating the surface expression levels of MHC II and CD86. PBMC were pre-treated with AOB, stimulated with the Th2 stimulus (by Th2 differentiation kit) for 72 hours, then immunostained with fluorescently labeled antibodies against the dendritic cell marker CD11c and either a) MHC II or b) CD86 followed by flow cytometry analysis. MHC II and CD86 expression were significantly reduced in CD11c+ cells after AOB treatment in 2 of 3 donors (Figures 3a-d). These observations suggest AOB can interfere with dendritic cell activation, and subsequent antigen presentation to T cells.

**Figure 3:**
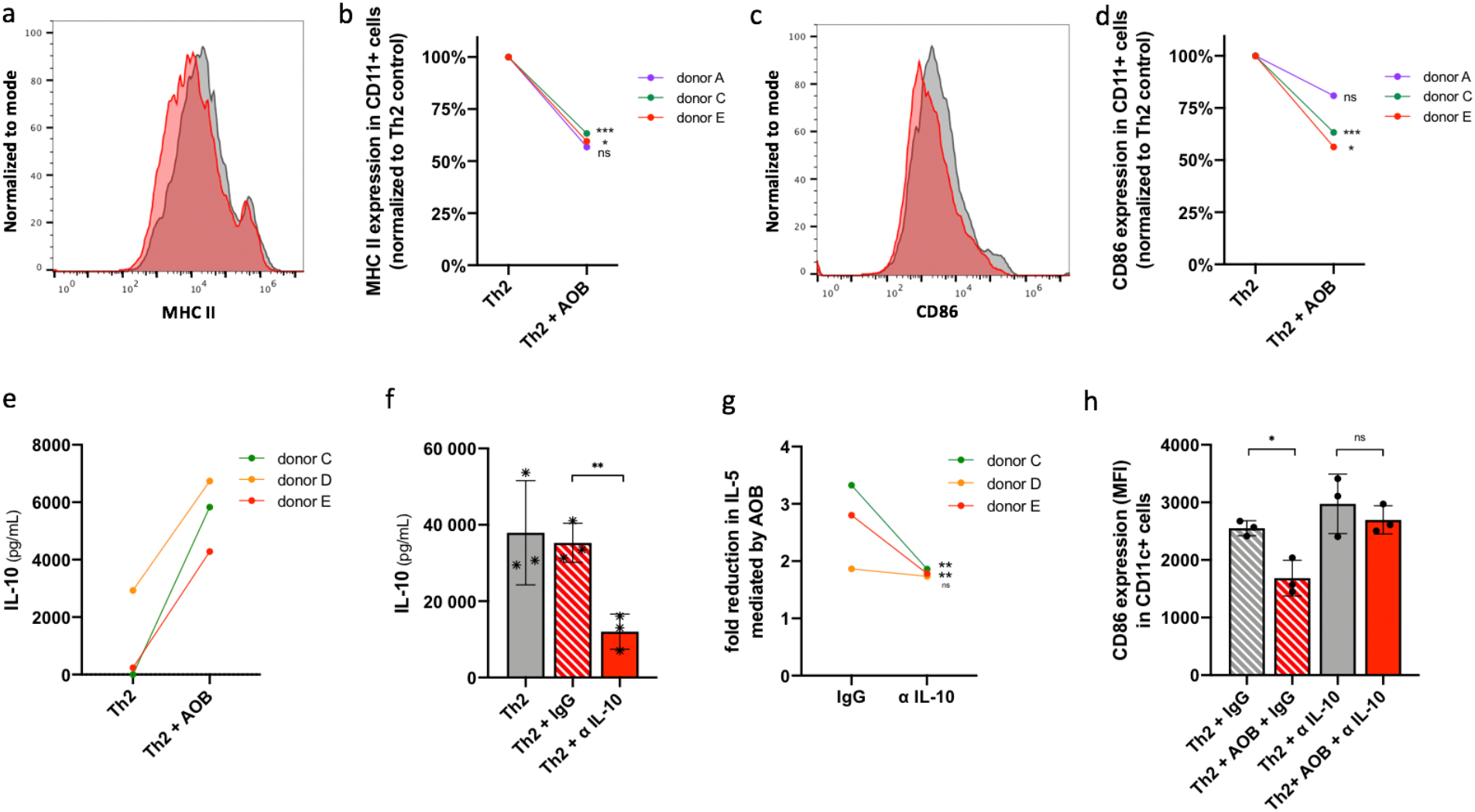
AOB-mediated Th2 suppression is IL-10 – dependent and involves a reduction of dendritic cell activation. (a-d) Expression of MHC II (a-b) or CD86 (c-d) in CD11c+ cells is reduced in PBMC treated with AOB prior to Th2 differentiation cocktail stimulation; measured by flow cytometry in Th2-stimulated PBMC in the presence or absence of AOB. (a, c) Representative fluorescence intensity plots of anti-MHC II FITC (a) or anti-CD86 PE (c) stained PBMC from donor C in the presence (red) or absence (grey) of AOB. (b, d) Relative MHC II (b) or CD86 (d) expression in CD11c+ positive cells is reduced with AOB treatment in 3 donors (n=3 per donor, unpaired t-test, p<0.0001 for MHC II and CD86 aggregated donor datasets). (e) IL-10 production is induced by AOB in PBMC from 3 donors 24h post-stimulation; measured by ELISA of culture supernatants (n=3 per donor, p=0.1095 for aggregated donor data set). (f-h) Addition of IL-10 neutralizing antibodies to PBMC prior to AOB pretreatment interfered with AOB’s inhibition of Th2. IL-10 (f) production by Th2-stimulated PBMC was reduced with the addition of IL-10 neutralizing antibody but not with isotype control in 3 donors (n=3 per donor, one point per donor, one-way ANOVA with multiple comparisons). AOB-mediated fold reduction in IL-5 (g) was obstructed by the addition of IL-10 neutralizing antibody compared to isotype control in 2 of 3 donors tested (n=3 per donor, unpaired t test). AOB-mediated reduction in CD86 expression (h) in CD11c+ cells is hindered with the addition of IL-10 neutralizing antibody but not with isotype control (n=3, donor D, one-way ANOVA with multiple comparisons).

The anti-inflammatory cytokine IL-10 can interfere with MHC II antigen presentation and CD86 co-stimulation on dendritic cells ^38,39^. We thus sought to test if AOB modulate dendritic cell activation via an IL-10 mediated mechanism. We first measured IL-10 production following AOB treatment and Th2 stimulation. We observed a potentiation of IL-10 production following AOB treatment and Th2 stimulation in all three donors tested (Figure 3e). Then, we tested if IL-10 was necessary for AOB-mediated inhibition of dendritic cell activity and the subsequent Th2 inhibition. IL-10 neutralizing antibody was applied prior to AOB treatment and Th2 stimulation. We verified that the anti-IL-10 antibody significantly reduced the levels of free IL-10 in the supernatant compared to the isotype control (Figure 3f, p<0.01). Interestingly, IL-10 neutralization significantly reduced AOB-mediated IL-5 inhibition in two out of the three donors tested (Figure 3g, p<0.01). We note that the donor D, which showed little response to the IL-10 antibody application, also showed the weakest inhibition of Th2 cytokines in response to AOB application following Th2 stimulation (Figure 1 d-e). Correspondingly, we observed that IL-10 neutralization also mitigated the AOB-induced suppression of CD86 expression in CD11c+ dendritic cells (Figure 3h). Together, these data support the concept that IL-10 plays an important, though not exclusive, role in AOB-mediated Th2 inhibition, most likely by interfering with dendritic cell ability to activate T cells.

### AOB – mediated Th2 inhibition doesn’t involve secreted bacterial metabolites

Next, we aimed to identify the bacterial component(s) responsible for Th2 inhibition. To discriminate between secreted metabolites and cell wall/intracellular components, we compared live AOB to AOB that were washed to remove any residual secreted molecules then heat killed to prevent any further metabolic activity. We verified that heat killed AOB were unable to produce nitrite or nitric oxide, indicating their lack of metabolic activity (Figures 4a-b, p<0.001). However, heat killed AOB were equivalent to live AOB in inhibiting the Th2 markers IL-5 and IL-13 (Figures 4c-d) or MHC II and CD86 expression (Figures 4e-f).

**Figure 4:**
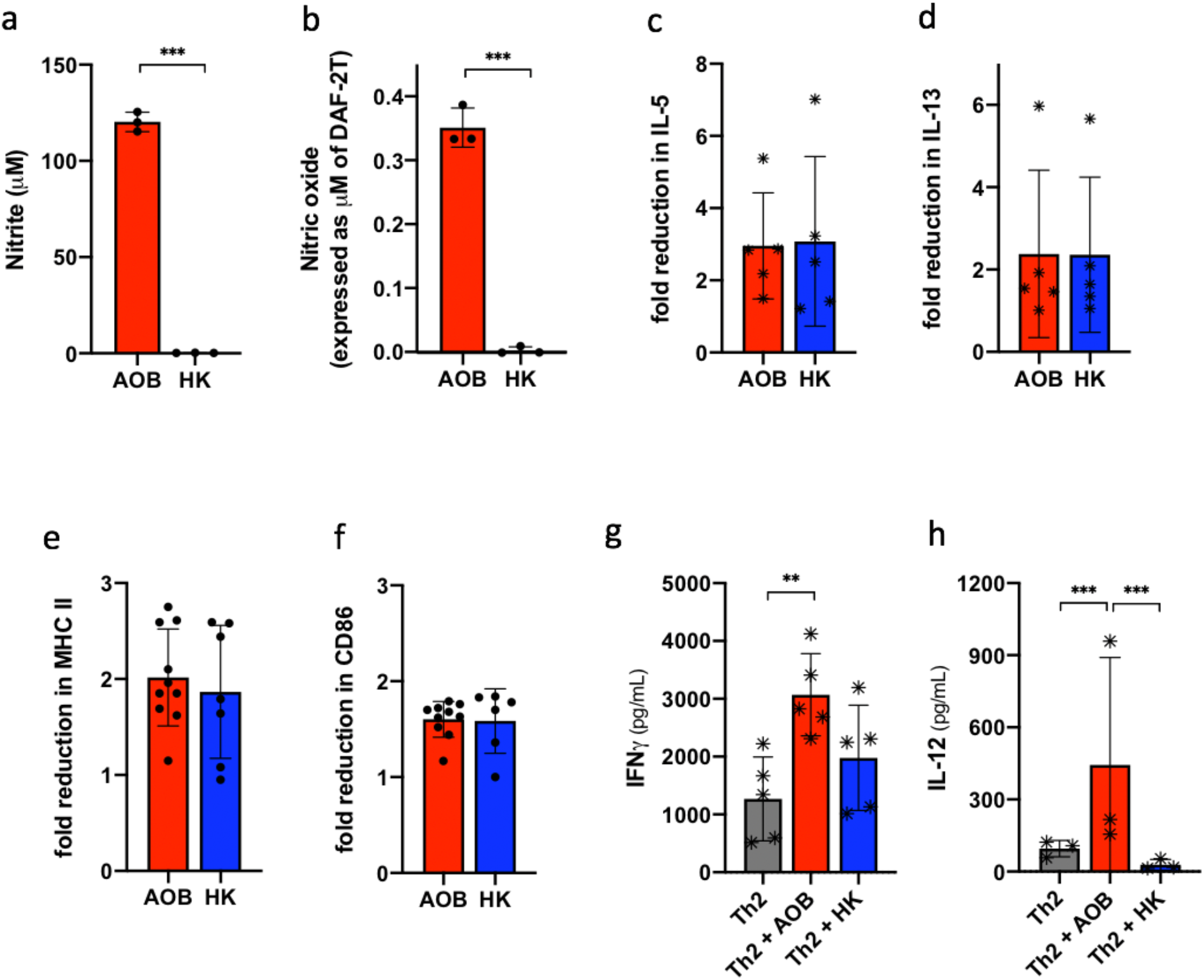
Structural components could be responsible for AOB-mediated Th2 inhibition. (a, b) AOB metabolites nitrite (a) and nitric oxide (b) are produced by live but not heat killed AOB after one hour incubation in AOB culture media (n=3, unpaired t-test). (c, d) Fold-reduction in IL-5 (c) and IL-13 (d) mediated by AOB (red) is not significantly different from fold-reduction mediated by heat killed AOB (blue); measured by ELISA in cell culture supernatant collected 72h post-stimulation with the Th2 stimulation cocktail (n**≥**6 per donor, 5 donors, one datapoint per donor, unpaired T test). (e, f) Fold reduction in MHC II (e) and CD86 (f) expression in CD11c+ cells mediated by live AOB (red) is not significantly different from fold-reduction mediated by heat killed AOB (blue); measured by flow cytometry 72h after Th2 stimulation in donor C (n=6, unpaired T test). (g, h) IFNγ (g) and IL-12p70 (h) production by PBMC are not as strongly induced by heat killed AOB (blue) as by live AOB (red); measured by ELISA from supernatant of PBMC culture collected 72h post-stimulation (n**≥**3 per donor, 3 or 5 donors, one datapoint per donor, one-way ANOVA with multiple comparisons).

Interestingly, the study of heat killed AOB further supports the differential role of Th1 polarization and dendritic cell activation. While heat killed AOB were equivalent to live AOB in regulating IL-5, MHC II, and CD86 expression (Figures 4c, e-f), heat killed AOB induced weaker levels of expression of IFNγ and IL-12 (Figure 4g-h).

In summary, AOB metabolic activity and associated secreted molecules, such as the immunomodulatory molecule nitric oxide, are not required for Th2 inhibition. Rather, these findings point towards a role of structural or intracellular components in mediating the effects on dendritic cells and subsequent Th2 polarization.

## DISCUSSION

This study reports the immuno-modulatory properties of the ammonia oxidizing bacteria *N. eutropha* D23. We discovered that AOB have the ability to suppress the polarization of CD4+ T cells to a Th2 phenotype by blocking dendritic cell activation in an IL-10 dependent manner. As such, this study highlights the potential for adding AOB to human skin to treat Th2-mediated pathologies associated with atopy. To our knowledge, this is the first report describing the immuno-modulatory properties of AOB.

### Mechanism of AOB-mediated Th2 modulation

The results presented here point to a non-canonical mechanism by which AOB suppress Th2 polarization in a Th1 independent manner, similarly to reports of a few other bacteria ^35,40^. While AOB can induce Th1 associated cytokines, we observed these cytokines are not a major contributor for Th2 suppression; IFNγ and IL-12 neutralizing antibodies did not prevent suppression, and suppression occurred with heat killed AOBs, which induced a weak IFNγ response.

Our data places dendritic cell modulation at the center of AOB’s mode of action. AOB reduced the surface levels of the proteins MHC II and CD86 involved in antigen presentation, and CD4+ T cell co-stimulation in CD11c+ dendritic cells, as shown with other immunomodulatory bacteria ^41^. Moreover, we found the anti-inflammatory cytokine IL-10 to be partially responsible for Th2 suppression, as has been described for other bacteria ^42^. We observed that IL-10 levels were increased with exposure to AOB, and more importantly, IL-10 neutralization using blocking antibodies prevented the AOB-mediated dendritic cell attenuation and the resulting Th2 suppression, an activity previously attributed to IL-10 ^38,39^. The origin of IL-10 induced by AOB, however, remains to be determined. Tolerogenic dendritic cells and regulatory T cells (Tregs) are common IL-10 producers in the context of immune tolerance ^43^. Tregs are unlikely to be involved, as AOB didn’t induce the Treg effector TGF-ß (*data not shown*). Moreover, the fact that AOB induce IL-10 as early as 24h suggests production by innate immune cells. Future studies that are beyond the scope of this paper will further address this point.

Unlike the Th1- and Treg-mediated immunomodulatory mechanisms of probiotics and gut commensals ^6,17,44^, environmental microbes such as cowshed bacteria and AOB appear to provide an alternate, additive source of signaling that is instructive to the immune system. We suggest that this alternate, instructive signaling can have important influences on the development of atopy ^45^.

### Role of metabolites and immuno-modulatory components

Exposure to environmental microbe components has long been associated with decreased prevalence of atopic diseases ^6,7,45^. Multiple types of bacterial immuno-modulatory molecules have been identified, from metabolites (e.g. SCFAs, indoles) to cell wall constituents (e.g. LPS, PSA) and nucleic acids (e.g. CpG DNA, RNA) ^6,46^. AOB’s ability to block Th2 polarization doesn’t seem to rely on metabolites, as heat killed AOB appeared as effective as live AOB. Therefore, the immuno-modulatory molecule nitric oxide produced by AOB is unlikely to be involved in this phenotype. We suspect that TLRs are involved in the AOB mechanism of action; future studies will test the relative activity of AOB on TLRs and aim to identify the bacterial substrates that imbue AOB with their immunomodulatory activity(s).

### Clinical utility of AOB

Atopic diseases such as atopic dermatitis, allergic rhinitis, and asthma involve an uncontrolled type 2 inflammatory response mediated by the Th2 cytokines IL-5, IL-13, & IL-4. IL-13 and IL-4 lead to the production of IgE and other mediators responsible for hypersensitivity reactions, tissue damage, and inflammatory reactions. IL-13 and IL-4 also bind to specific receptors at the surface of sensory neurons and play a role in promoting itch ^47^. IL-5 is involved in eosinophil activation, effectors of which lead to tissue damage ^48,49^. Previous approaches suggested that the efficacy of targeting end mediators/effectors was not as widespread across atopic diseases as blocking upstream Th2 cytokines (e.g. IL-5, IL-13, IL-4) ^14^. Based on the ability of AOB to modulate all three major Th2 cytokines, we hypothesize that AOB have the potential to influence atopic disease by blocking both IgE and eosinophil pathways, as well as interfering with itch signaling. However, future studies will be required to determine if AOB indeed have this activity.

Potential limitations to this study include the reliance on Th2 stimuli that do not fully mimic the diseased state, as well as the use of immune cells with which an environmental microbe would not frequently come into close contact. However, our observation that AOB suppress Th2 polarization mediated by Staphylococcal enterotoxin B (SEB) provides confidence that our results are not due to spurious activity of a proprietary Th2 stimulation cocktail. Furthermore, and more importantly, the SEB observations reinforce AOB’s clinical potential due to the relevance of *Staphylococcus aureus* and SEB in atopic dermatitis. In this disease, one might expect dermally applied AOB to encounter immune cells in a manner similar to our study, as patients often breach the skin barrier in response to the intense itch associated with SEB. Future preclinical and clinical studies will provide more insight on whether AOB-mediated Th2 inhibition can translate into disease modifying phenotypes in atopic disease.

Heat killed AOB, which retain immuno-modulatory activity, might be leveraged to prevent colonization and potential adverse events commonly associated with live bacterial products ^50^. Moreover, their lower Th1 induction could be beneficial in the context of atopic diseases with a Th1 component (e.g. celiac disease) or atopic patients also suffering from Th1-mediated diseases (e.g. multiple sclerosis, type 1 diabetes) ^51^.

In conclusion, the ammonia oxidizing bacteria *N. eutropha* D23 shows promising therapeutic potential to target atopic diseases due to its ability to block Th2 polarization and key cytokines involved in IgE production, eosinophilia, & itch. Follow up studies will provide more insights into its detailed mode of action and clinical potential.

## METHODS

### Bacteria and PBMC

*Nitrosomonas eutropha* D23 is a unique strain of ammonia-oxidizing bacteria (AOB) manufactured by AOBiome Therapeutics. AOB were washed in PBS (Gibco) prior to being used for PBMC treatment. Heat killed AOB were prepared by first washing live AOB in PBS and then incubating AOB at 60°C for 2 hours.

Human PBMC were purchased from ATCC. 5 different PBMC donors were used with the following lot numbers from ATCC: 80227190 (donor A), 80424333 (donor B), 80819190 (donor C), 80424222 (donor D), and 80808190 (donor E).

### PBMC stimulation and treatments

PBMC were seeded at a concentration of 7-8 × 10^6^ cells/mL in RPMI 1640 medium without phenol red and glutamine (Gibco) supplemented with 10% heat-inactivated FBS (Gibco) and 2mM glutamax (Gibco) and incubated at 37°C with 5% CO_2_ overnight. AOB were added at a ratio of four bacteria per PBMC one hour prior to Th2 stimulation. PBMC were then stimulated using either the CellXvivo human Th2 differentiation kit (R&D systems) following the manufacturer’s recommendations, or 1μg/mL SEB (Sigma). Where appropriate, neutralizing antibodies were used at 10μg/mL and refreshed daily until sample collection. The following neutralizing antibodies were used: mouse anti-human IFNγ clone K3.53 (MAB2852), mouse anti-human IL-12p70 clone 24910 (MAB219), mouse anti-human IL-10 clone 948505 (MAB9184) and the isotype controls mouse IgG2a (MAB003) and mouse IgG1 (MAB002) (all from R&D systems).

Samples were collected three days post-stimulation (unless otherwise indicated) by spinning down cells at 500 x g for 10 min. Cell-free supernatants were frozen at −20°C for later cytokine quantification and cell pellets were processed for PBMC viability, RNA extraction or flow cytometry analyses as described below.

### PBMC viability

PBMC viability was determined by measuring cell metabolic activity using WST-1 as described by Klinder et al.^52^. Briefly, 50μl aliquots of cells were transferred into a 96 well plate, diluted 2-fold with PBMC culture media and supplemented with 10% WST-1 reagent (Roche) according to manufacturer’s instructions. Samples were incubated for 1 hour at 37°C with 5% CO_2_ and OD_440nm_ & OD_650nm_ were then measured using a 96-well plate spectrophotometer (Molecular Devices or BioTek). OD_650nm_ was subtracted from OD_440nm_ then percent viability was calculated relative to the no treatment condition.

PBMC viability was also confirmed using LIVE/DEAD flow cytometry staining. 1×10^6^ cells were washed in twice in PBS and stained with the LIVE/DEAD Fixable Green Dead Cell Stain kit for 488 nm excitation (Invitrogen) following the manufacturer’s instructions. Cells were incubated in the dark for 30 minutes, washed twice in PBS + 1% BSA then analyzed using a BD Accuri C6 flow cytometer. Data were quantified using FlowJo 10 software.

### Cytokine quantification by ELISA

Supernatant samples were diluted in PBMC culture media prior to being assayed in order to be in the ELISAs’ dynamic range. ELISAs were then performed following manufacturer’s instructions and cytokine concentrations were calculated based on a linear regression of the standard curve. IL-5, IL-13 and IL-12 p70 ELISA kits were purchased from R&D systems, IFNγ and IL-10 from BD Biosciences).

### Flow cytometry analysis of surface and intracellular markers

For intracellular quantification of IFNγ or IL-5, cells were treated with 4μM Golgistop (BD Bioscience) to block protein secretion 14 hours prior to sample collection. In the case of IL-5, cells were also treated with 81nM PMA and 1.3μM ionomycin (eBioscience) 5 hours prior to sample collection to increase cytokine detection threshold. Cells were then washed twice in FBS buffer (BD Bioscience), fixed in BD cytofix (BD Bioscience), washed twice in FBS buffer and frozen at −80°C in 90% heat inactivated FBS – 10% DMSO until further analysis. Thawed cells were then washed twice in FBS buffer and permeabilized with BD perm/wash buffer (BD Bioscience). For IL-5 staining, cells were blocked with 4μg of unlabeled rat IgG1 clone R3-34 antibody (BD bioscience). 1×10^6^ cells were then stained with the following antibodies: mouse anti-human CD4 PerCP clone SK3 antibody (20μL), mouse anti-human IFNγ FITC clone B27 (20μL), rat anti-human IL-5 PE clone TRFK5 (5μL) and their respective isotypes mouse IgG1 FITC clone MOPC-21 (20μL) and rat IgG1 PE clone R3-34 (5μL) (all from BD Bioscience). Stained cells were then washed twice in perm/wash buffer then resuspended in FBS buffer and analyzed on a BD Acuri C6 flow cytometer (BD Bioscience), and data were quantified using FlowJo 10 software.

For quantification of the surface markers MHC II and CD86, cells were washed twice with FBS buffer. 1×10^6^ cells were then stained with the following antibodies: mouse anti-human MHC II FITC clone Tu39 (20μL), mouse anti-human CD86 PE clone 2331 (FUN-1) (20μL), mouse CD11c APC clone B-ly6 (20μL) and their respective isotypes mouse IgG2a FITC clone G155-178 (20μL), mouse IgG1 PE clone MOPC-21 (20μL) and mouse IgG1 APC clone MOPC-21 (all from BD Bioscience). Stained cells were then washed twice in FBS buffer then resuspended in FBS buffer and analyzed on a BD Acuri C6 flow cytometer (BD Bioscience), data were quantified using FlowJo 10 software.

### Gene expression analysis by RTqPCR

Due to the presence of high levels of IL-4 in the Th2 differentiation kit preventing a robust estimate of AOB impact on IL-4 production, IL-4 production was measured by RTqPCR to selectively quantify cellular IL-4. PBMC were washed in PBS then subjected to RNA extraction using the Qiagen RNAeasy extraction kit including the QIAshredder homogenization step following manufacturer’s recommendations. RNA was then reverse-transcribed into cDNA using the Invitrogen SuperScript III First strand synthesis system for RTqPCR following the manufacturer’s protocol with oligo dT primers. Gene expression was then measured by qPCR using the PowerUp SYBR Green Master Mix (Applied Biosystems) and QuantStudio 3 or 6 Flex (ThermoFisher Scientific) following manufacturer’s recommendations. qPCR reactions were performed in a final volume of 25μL with an annealing temperature of 55°C for 40 cycles. The following primer sequences were used: IL-4 (F: CCAACTGCTTCCCCCTCTG, R: TCTGTTACGGTCAACTCGGTG; sequences from PrimerBank ^53^), EF-1a (F: CTGAACCATCCAFFCCAAAT, R: GCCGTGTGGCAATCCAAT ^54^. Relative gene expression was calculated using the 2^-ddCt method ^55^ using EF-1a as a housekeeping gene ^54^.

### Bacterial metabolite quantification

Nitrite and nitric oxide production were measured to determine the metabolic activity of live or heat killed AOB. Nitrite was quantified using the Griess assay. Briefly, samples were mixed 1:1 with a solution of equal volumes Griess Reagent A (1.5 N Hydrochloric Acid, 58 mM Sulfanilamide) and Griess Reagent B (0.77mM NNEQ), incubated for 20 minutes at room temperature protected from light then quantified by measuring OD_540nm_ using a 96 well plate spectrophotometer. Concentrations of nitrite were then calculated based on a standard curve of sodium nitrite. Nitric oxide was quantified using the fluorescent dye DAF-2 (Abcam). Briefly, samples were mixed 1:1 with PBMC/AOB culture media containing 5μM DAF-2 and incubated at 37°C with 5% CO_2_ for 1 hour. A standard curve of the fluorescent molecule DAF-2T (Abcam) was used to quantify nitric oxide production. Measurements were either taken after one hour in AOB culture media or 72 hours in PBMC culture media.

### Statistical analysis

As bacterial interventions affect host physiology based on the variable genetic makeup of the host, we have chosen to analyze comparisons within donors, as opposed to across donors. The numbers of donors tested in this study would not offer sufficient statistical power to make proper inferences; clinical studies often require patient numbers in the 100s, even when measuring clinical biomarker samples using ELISAs following use of drugs with a known mechanism of action. One-way analysis of variance (ANOVA) with Tukey multiple comparison correction or unpaired t-test were performed when appropriate, as indicated in the figure legends, using Graphpad Prism 8 software (Graphpad Software Inc).

## ACKNOWLEDGMENTS

We would like to thank AOBiome Therapeutics for providing the funding and resources for this study as well as all AOBiome’s team members, particularly Daniel Brownell, Judith Ng-Cashin, Ioannis Gryllos, Chris Johnson, and Neeraja Vajrala. Additionally, we would like to acknowledge Robrecht Thoonen and Anita Hartog for their advice and review of the manuscript.

## CONFLICT OF INTEREST

The authors were employed by AOBiome Therapeutics at the time of this study; AOBiome Therapeutics is currently investigating the clinical use of AOB for treatment of atopic dermatitis.

## AUTHOR CONTRIBUTIONS

AG and DM contributed to the design and concept; DM and NE conducted the experiments; all authors contributed to results analysis, interpretation, and final manuscript preparation.

