## Supplementary figures for "The ammonia oxidizing bacterium *Nitrosomonas eutropha* D23 blocks T helper 2 cell polarization via IL-10 – mediated interference with dendritic cell activation"

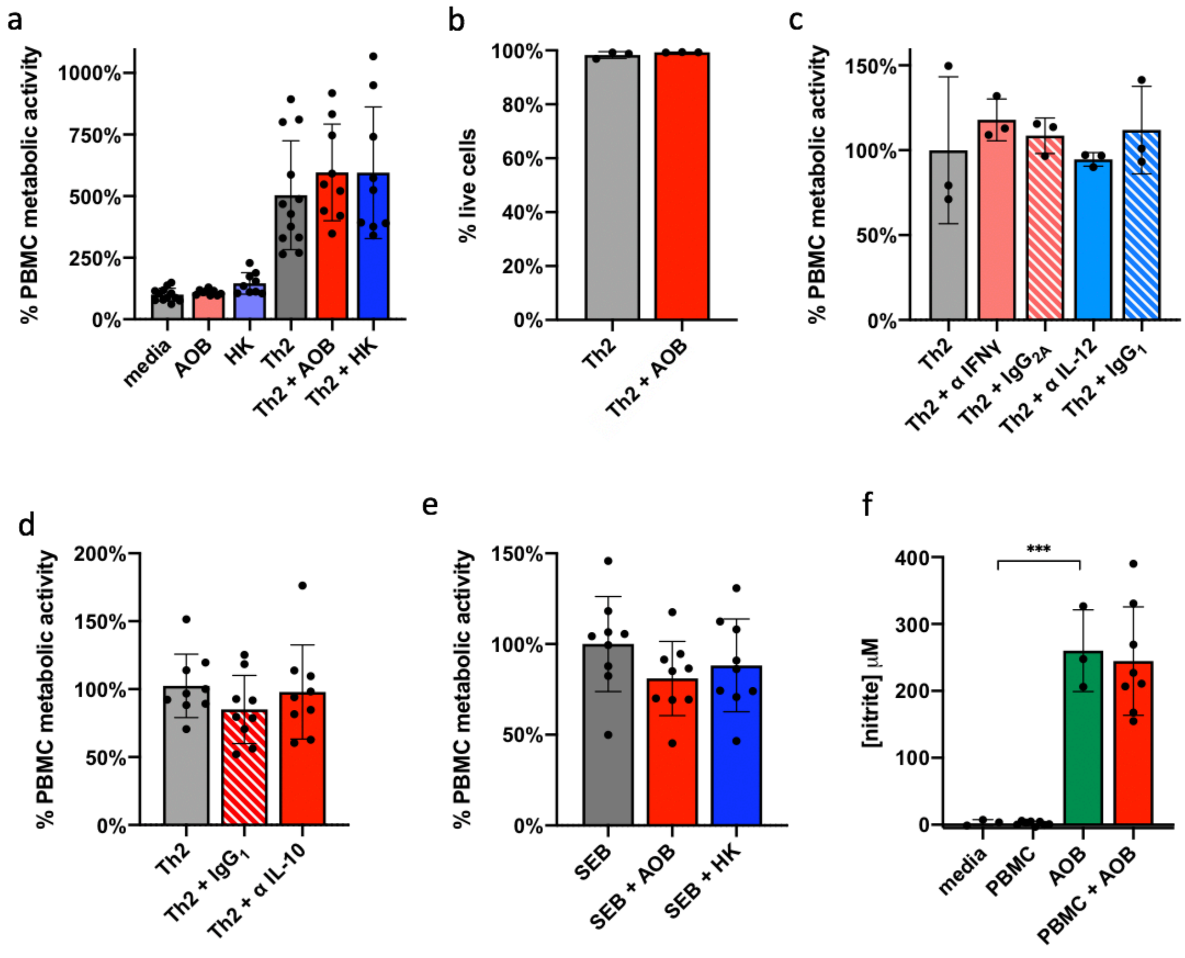


**Figure S1 – Various AOB or blocking antibody treatments are not toxic to PBMC, and AOB retain metabolic activity when in contact with PBMC**

(a) PBMC metabolic activity is unaffected by addition of live or heat killed AOB and increased after Th2 stimulation; measured by WST-1 as an indicator of cell viability in samples collected 72h after treatment (n≥9, donor A, one-way ANOVA with multiple comparisons). (b) Percentage of live PBMC is unchanged in presence or absence of AOB; measured by LIVE/DEAD staining (n=3, donor C, unpaired t test). (c) PBMC metabolic activity is not significantly different in the presence or absence of IFNɣ or IL-12 neutralizing antibodies or isotype controls; measured by WST-1 (n=3, donor A, one-way ANOVA with multiple comparisons) (d) PBMC metabolic activity is not significantly different in the presence or absence of IL-10 neutralizing antibody or isotype control; measured by WST-1 (n=3 per donor, 3 donors, one-way ANOVA with multiple comparisons). (e) PBMC metabolic activity after SEB stimulus is not significantly different in presence or absence of live or heat killed AOB (n=3 per donor, 3 donors, one-way ANOVA with multiple comparisons). (f) Nitrite is not produced by PBMC alone but is produced by AOB both in presence and absence of PBMC; measured by Griess assay (n≥3, one-way ANOVA with multiple comparisons).

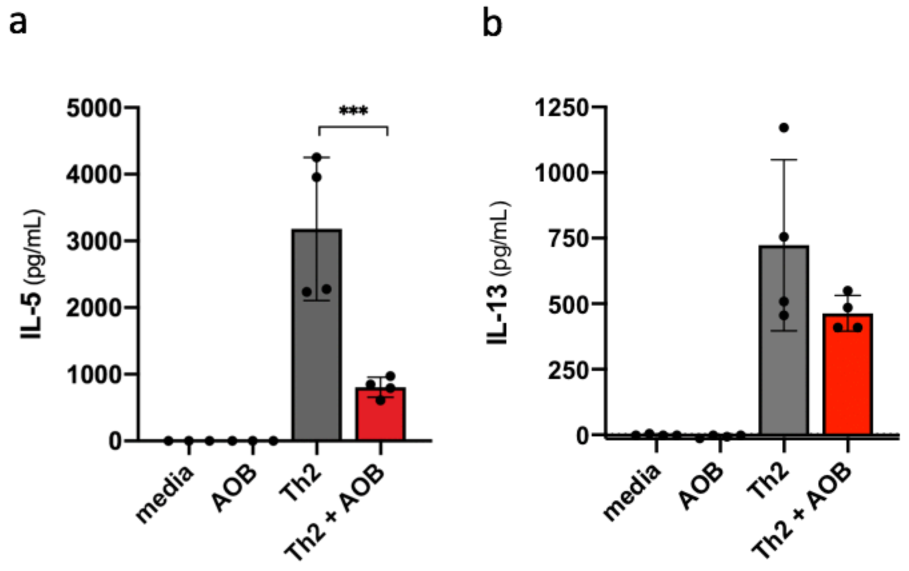


**Figure S2 – AOB reduce the production of Th2 cytokines by PBMC collected 5 days post-stimulation**

(a, b) IL-5 (a) and IL-13 (b) production is reduced with AOB pretreatment prior to stimulation by Th2 differentiation cocktail; measured by ELISA from culture supernatants of PBMC collected 5d post-stimulation (n=4, donor C, one-way ANOVA with multiple comparisons).
